# EpiRank: partner-agnostic surface-patch retrieval for epitope prediction of protein binding interfaces

**DOI:** 10.64898/2026.10.26.762091

**Authors:** Brian Lin, Dongsheng Lu

## Abstract

Structure-based interface predictors are normally reported as a single pooled ROC AUC over a residue test set. For a protein designer the operative question is different and much harder: *given one target chain, will the top-ranked surface patch be a real binding site?* We built EpiRank, a partner-agnostic epitope predictor, on HumanPPI — 29,246 predicted human dimers, 12,296 target chains, 4.18 M residues — using only single-chain geometry and physicochemistry (170 features, no partner, no sequence search, no structural alphabet), and evaluated it as a retrieval problem with homology- and partner-graph-controlled splits, on the 3,529 chains that survive an explicit partner-contamination filter. The model places a real epitope in its top-ranked patch 64 % of the time. That number is far less informative than it looks. Compact patches placed at random hit a real epitope 41 % of the time under the same rule, so the model’s lift is 1.57×, not 64 %; 54 % of all predicted patches land on nothing; and when the same chains are scored by models trained on a 2,000-chain and an 11,669-chain superset, a substantial fraction change answer while the aggregate barely moves. We document four specific ways this task invites overstatement — partner-graph percolation across splits, single-linkage homology percolation at the conventional 30 % identity threshold, a count-only hit rule that rewards spatial smoothing rather than signal, and hotspot precision conditioned on success, which ranks a random baseline *above* the trained model. Each was hit during development and each inverted a conclusion we had already drawn. We argue that interface-prediction papers should report lift over a geometry-matched random baseline, patch precision, and per-target stability alongside AUC, and we release the evaluation harness that computes them.

## 1. Introduction

Designing a minibinder or a VHH against a new target begins with a choice that is made before any design software runs: *which patch of the surface to aim at*. Current generative pipelines — RFdiffusion [1], BindCraft [2], BoltzGen — take hotspot residues as an input, not an output. A wrong hotspot set does not produce a worse binder; it produces a confident binder against a site that does not matter, and the cost is an entire wet-lab campaign.

Structure-based interface prediction should answer this. The field has matured quickly: MaSIF learned surface fingerprints from geometry and chemistry [3], ScanNet learned interpretable atom-to-residue filters end to end [4], and PeSTo removed hand-built features altogether with a parameter-free geometric transformer [5]. Reported performance is high — PeSTo reports residue-level ROC AUC close to 0.9 on protein–protein interfaces [5]. Separately, Foldseek-Interface [6], built on the Foldseek structural alphabet [7], showed that the interface universe sampled by the PDB is small and strongly redundant, which reframes what a held-out test set actually holds out. The surface-patch framing itself is much older [8], as is the observation that interface residues occupy a characteristic structural region of the surface [9].

We set out to build a predictor that was deliberately modest in its assumptions, for a specific reason: we wanted to know what a designer actually gets from a surface-feature model, measured the way a designer would use it. That led to three departures from the standard evaluation.

### The label is partner-agnostic

We do not predict “the interface with partner X”. We take the union of contacts across every partner a chain has in the dataset, so a positive residue means *some* protein binds here. This is the right target for binder design — the designer does not care which natural partner defines the site — but it means a chain observed with one partner is labelled incomplete, and the negatives mean “no partner was modelled”, not “does not bind”.

### The unit of evaluation is the protein, not the residue

Pooled residue AUC weights a 900-residue chain 6× more than a 150-residue one, and tells the designer nothing about whether *their* target will work. We score each chain independently and report the distribution.

### The prediction is a ranked list of patches

A designer takes the top one, maybe the top three. So we ask: does the top-ranked compact patch overlap a real epitope, and are its highest-scoring residues real contacts?

The result of measuring this way is that a model with an unremarkable AUC turns out to be genuinely useful — 1.57× over random on the metric that matters, and 1.74× on the hotspot residues a designer actually uses — *and* that almost every intuitive way of reporting it inflates the number. Section 4 is devoted to the inflation mechanisms, because we believe they are general to the task rather than specific to our model.

## 2. Results

### 2.1 A partner-agnostic benchmark from predicted human dimers

HumanPPI provides AlphaFold2 [10], AlphaFold-Multimer [11] and RoseTTAFold [12] models of 29,246 human protein pairs, part of the broader effort to resolve the human interaction network structurally [13,14]. We extracted 12,296 target chains in isolation and labelled a residue positive if any heavy atom lies within 5.0 Å of any heavy atom of a partner chain in any complex that chain participates in (Figure 1a). All results below are on the complete set unless stated otherwise.

**Figure 1.**
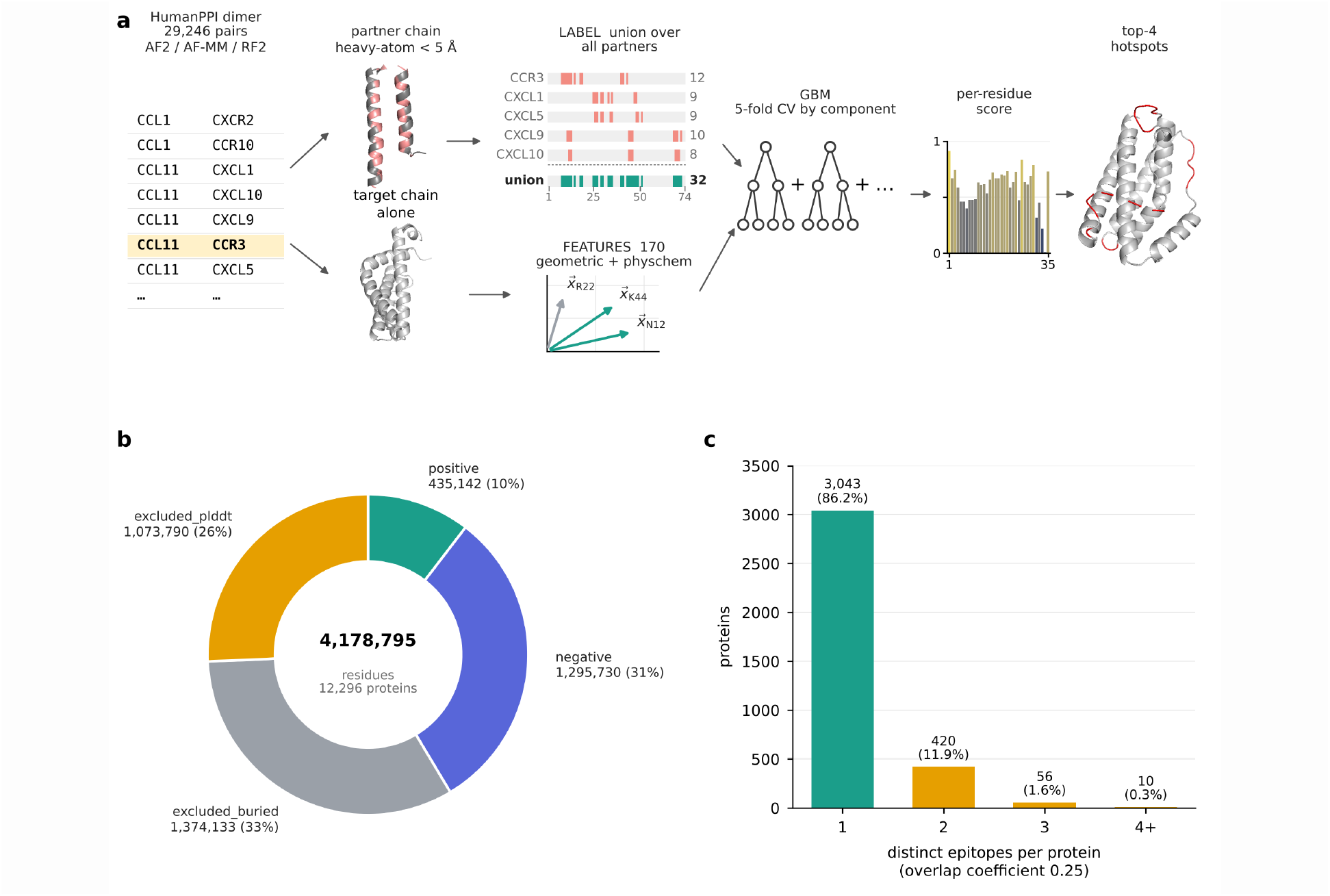
Benchmark construction. (a) Pipeline — the partner chain is used for labels only, never for features. CCL11 is shown as a worked example: its contacts with five partners (CCR3, CXCL1, CXCL5, CXCL9, CXCL10) union to a 32-residue label over a 74-residue chain, and features are computed on the unbound chain alone. (b) Residue classes over the 4,178,795 classified residues of 12,296 chains — buried and low-confidence residues are excluded from training and from every metric. (c) Of the 3,529 evaluated chains, 14 % carry more than one binding site; the training label unions them, the evaluation keeps them apart.

Two filters are applied *before* training and are applied identically in every metric reported here. Residues with relative solvent accessibility below 0.15 are buried and cannot be targeted; residues with mean pLDDT below 70 are not reliably placed. Together they remove 59 % of all residues (Figure 1b). This matters for interpreting the AUC: a model that merely separates buried from exposed residues scores well on an unfiltered set and is useless to a designer, because every candidate the designer could use is already exposed. On our filtered set the positive rate is 25 %.

25 % of target chains carry more than one spatially distinct binding site, defined by clustering the per-partner contact sets at an overlap coefficient of 0.25. A 2,000-chain pilot subset gave 42 % — the pilot was enriched for well-connected hubs, which is one more reason to report the complete set. The subset that survives the contamination filter of §2.5 is depleted of hubs further still, to 14 % (Figure 1c): a chain with many partners has more refused merges, so it is more likely to be excluded. The training label unions these sites; the evaluation keeps them apart, which is what makes hit@*k* and recall@*k* different questions.

**Table 1.** Benchmark composition.

| **Quantity** | **Value** |
|---|---|
| Source pairs in HumanPPI | 29,246 |
| Target chains | 12,296 |
| Residues total | 4,180,280 (4,178,795 classified) |
| Positive (interface) | 435,142 (10.4 %) |
| Negative (surface, no partner) | 1,295,730 (31.0 %) |
| Excluded, buried (RSA < 0.15) | 1,374,133 (32.9 %) |
| Excluded, low pLDDT (< 70) | 1,073,790 (25.7 %) |
| Unmodelled | 1,485 (< 0.1 %) |
| Positive rate on the evaluated set | 25 % |
| Chains with epitopes mapped | 9,342 |
| Chains with ≥ 2 distinct epitopes | 25 % |
| Partner-contaminated, excluded from metrics | 5,813 |
| **Chains in the headline evaluation** | **3,529** |
| ≥ 2 epitopes within the evaluated subset | 14 % |
| Independent components after split control | 1,345 (largest 679 chains) |
| Held out — never trained, never tuned | 1,698 chains (259,860 res., 15 %) |
| Cross-validation folds | 5 × ≈294,200 residues |

### 2.2 Features: single-chain geometry and physicochemistry only

170 features per residue, computed on the unbound chain: relative solvent accessibility; burial depth; local curvature and convexity from the Cα shell; rotation-invariant local-frame descriptors of the neighbourhood geometry; secondary-structure context; residue identity and standard physicochemical scales; and neighbour-averaged versions of all of the above at two radii.

We deliberately excluded three things. No multiple-sequence-alignment or conservation features, because they require a sequence search and because conservation partly encodes the answer. No structural-alphabet descriptor (Foldseek 3Di [7] or similar) — we use the underlying idea, that interface geometry is a discretisable local property, but compute it directly rather than through an alphabet trained for a different objective. And no partner-derived quantity of any kind, including the solvent accessibility of the complex.

That last exclusion is not a minor hygiene point. Our first model reached a pooled AUC of 0.999. The cause was `delta_rsa` — the change in accessibility on complexation — which had been carried into the feature table. It *is* the label, expressed as a float. We now compute RSA on the isolated chain only, and every complex-derived quantity lives in the label file.

### 2.3 Residue-level performance is unremarkable, and that is informative

A histogram-based gradient-boosting classifier trained with 5-fold cross-validation (splits described in §2.5) reaches a pooled ROC AUC of **0.719** and a per-chain median of **0.687** over the 3,572 uncontaminated chains that carry both classes (Figure 2). The precision–recall curve starts near 0.95 and decays to the 0.22 base rate.

**Figure 2.**
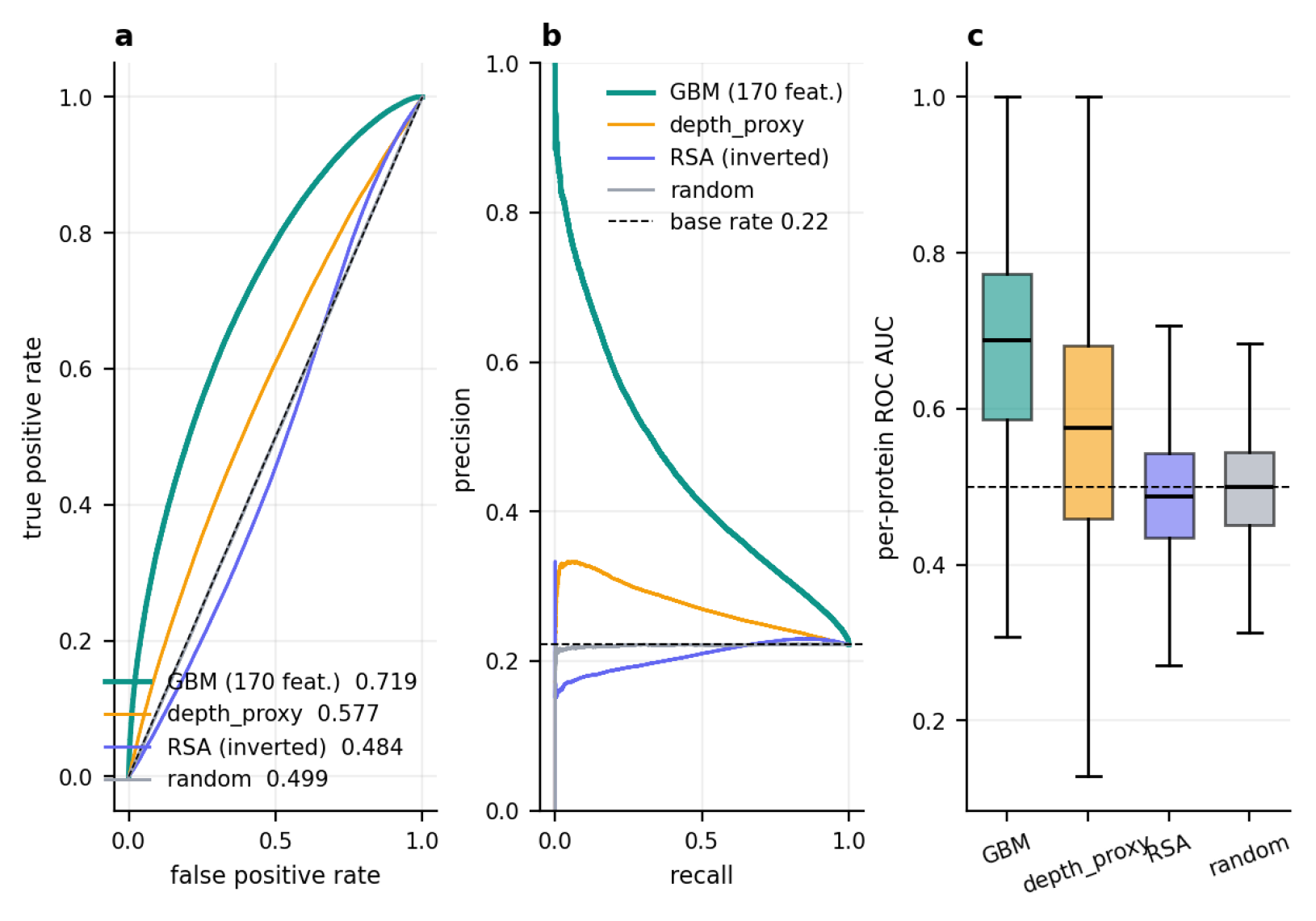
Residue-level performance on the uncontaminated subset. (a) Pooled ROC. (b) Precision–recall against the 0.22 base rate. (c) Per-chain AUC distribution over 3,572 chains. RSA alone is below chance once buried residues are excluded.

The baselines are the point of this figure. A simple burial-depth proxy reaches 0.577 pooled; inverted RSA reaches **0.484** — *below chance*. This is the direct consequence of the surface filter. In an unfiltered evaluation, “is it exposed?” is most of the signal, and a published claim we had carried in our own planning documents — that buried-surface features alone give ROC AUC around 0.9 — simply does not survive the filter. We measured 0.679 for the best single feature and 0.630 for RSA alone before filtering; after filtering RSA is anti-correlated with interface status, because the most exposed residues on a folded chain are disordered loops and termini, not binding sites.

An AUC of 0.72 is well below the ≈0.9 that PeSTo [5] and comparable models [3,4] report. We do not claim our features match theirs. But the gap is not purely a model gap: it is partly an evaluation gap. Our positives are partner-agnostic unions on predicted structures with an unknown fraction of unobserved partners mislabelled as negative, our buried residues — the easiest true negatives — are removed before scoring, and our splits are controlled for both homology and partner sharing. Each of those choices costs AUC and none of them can be reversed without making the benchmark less honest for the design use case.

### 2.4 Retrieval: the metric a designer should read

Per-residue scores are smoothed over a 10 Å Cα neighbourhood, the top 25 % are retained, and connected components within 8 Å form candidate patches, ranked by size × mean score (Figure 1a, lower row). A patch counts as a hit if it shares at least 3 residues with one epitope **and** at least 25 % of the patch lies inside that epitope. The second clause is essential and is justified in §4.3.

**Table 2.** Retrieval performance, 3,529 uncontaminated chains.

| **Metric** | **GBM** | **depth proxy** | **random** | **GBM lift** |
|---|---|---|---|---|
| hit@1 | **64 %** | 54 % | 41 % | **1.57×** |
| hit@3 | **79 %** | 67 % | 59 % | 1.33× |
| recall@5 (all sites found) | **77 %** | 64 % | 59 % | 1.30× |
| patch precision (pooled, 10,546 patches) | **46 %** | 41 % | 28 % | 1.64× |
| patch precision, top 3 | **56 %** | 49 % | 37 % | 1.51× |
| hotspot precision (top 4 residues of top patch) | **49 %** | 37 % | 28 % | **1.74×** |
| median ΔSASA of correct hotspots | 50 Å² | 43 Å² | 43 Å² | — |

Every absolute rate is lower than on the 2,000-chain pilot (hit@1 70 % → 64 %) and every lift is higher (1.35× → 1.57×; hotspot 1.53× → 1.74×). Two effects push in the same direction. The complete set holds more single-site chains and smaller epitopes, so the random baseline has less to hit by accident; and removing partner-contaminated chains (§2.5) takes a further 3.1 points off the model’s hit@1 while barely moving random’s. A larger and better-controlled benchmark made the headline rate worse and the measured signal stronger. That is the expected direction, and it is the argument for reporting lift rather than rate.

The random baseline is not a shuffled score. It is a compact patch of matched size placed at a random surface location, smoothed and clustered through the identical pipeline — the correct null for a spatial retrieval task. Against it, hit@1 of 64 % is a 1.57× lift, and hit@3 of 79 % is only 1.33×, because a chain with several epitopes is easy to hit by chance three times. **The headline number a designer should take from this work is the hotspot lift of 1.74×, not the hit rate of 64 %**.

Two further observations (Figure 3). Chains with more epitopes are easier to hit once and harder to cover completely: hit@1 rises monotonically from 63 % (one site, n = 3,043) to 70 % (two, n = 420) to 74 % (three or more, n = 66) and hit@3 from 77 % to 86 % to 91 %, while recall@5 falls from 79 % to 62 % to 55 %. Finding *a* site gets easier and finding *every* site gets harder, and the two trends are often reported as one number. And patch precision is bimodal — the model is either largely right or largely wrong on a given chain, with 54 % of all emitted patches landing on nothing. That 54 % is the wasted-design-run rate, and it is the number we would want on the first slide of any deployment discussion.

**Figure 3.**
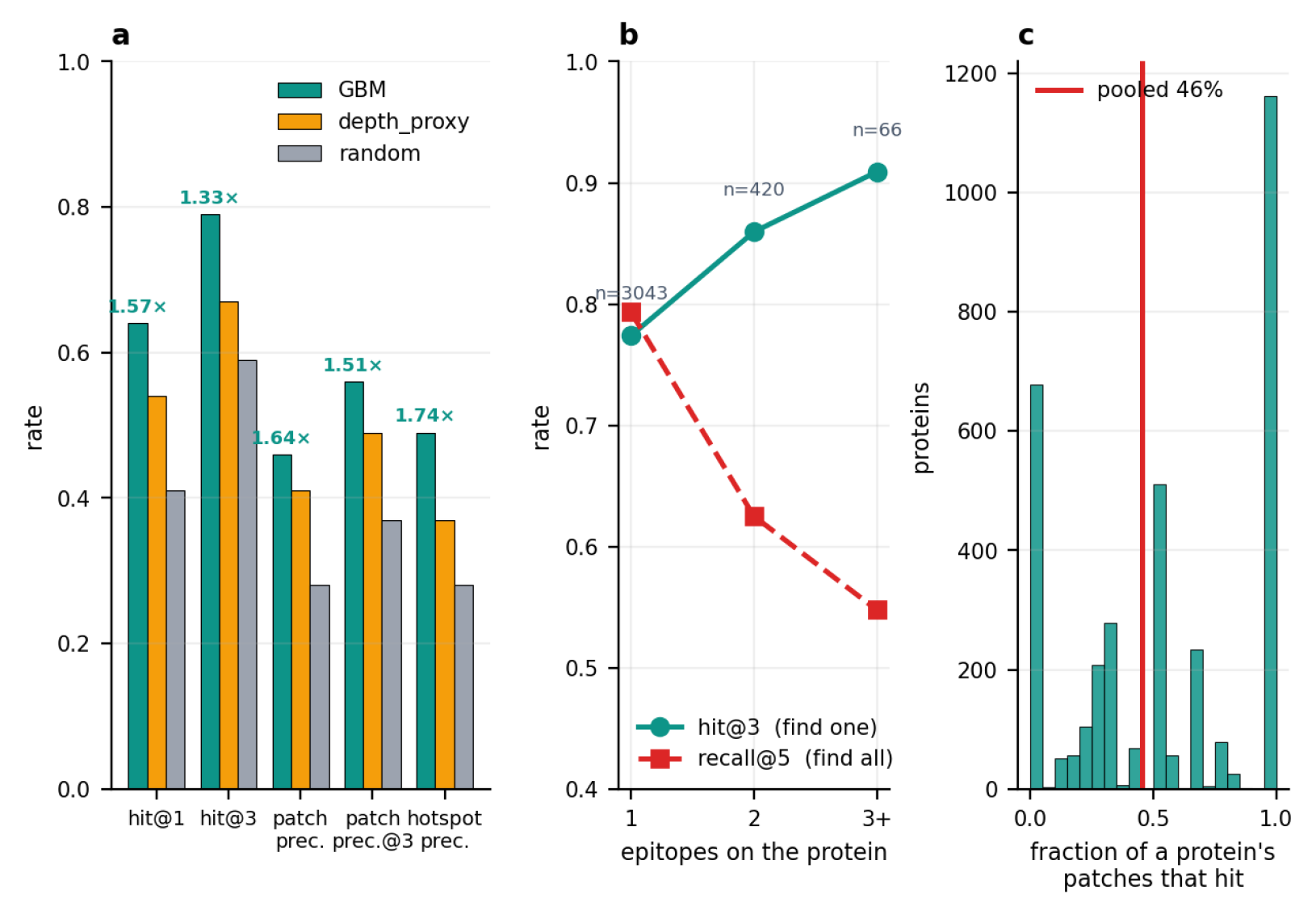
Retrieval performance over the 3,529 uncontaminated chains. (a) All metrics against both baselines, with lift annotated. (b) Multi-site chains: easier to find one site, harder to find all. (c) Per-chain patch precision is bimodal; 54 % of patches hit nothing. The random baseline is high because the hit rule is deliberately permissive — read the lift, not the rate.

### 2.5 Split construction: two percolation failures

Standard practice is to cluster by sequence identity at 30 % and split by cluster. Applied to the complete set, single-linkage clustering at exactly 30 % puts **97.1 % of all residues in one component** (Figure 4b). Not 97 % of chains in a few big families — one component, containing essentially the whole dataset. At 40 % it still holds 36.1 %, and only at 50 % does it fall to 5.4 %. Chains A–B and B–C each below 30 % identity to one another are merged through B, and with enough chains the transitive closure swallows everything. The 2,000-chain pilot gave 29.2 % at the same threshold, so this is a property that appears only at scale: **a homology-controlled split by single linkage at the field-standard threshold is not possible on this dataset at all**.

**Figure 4.**
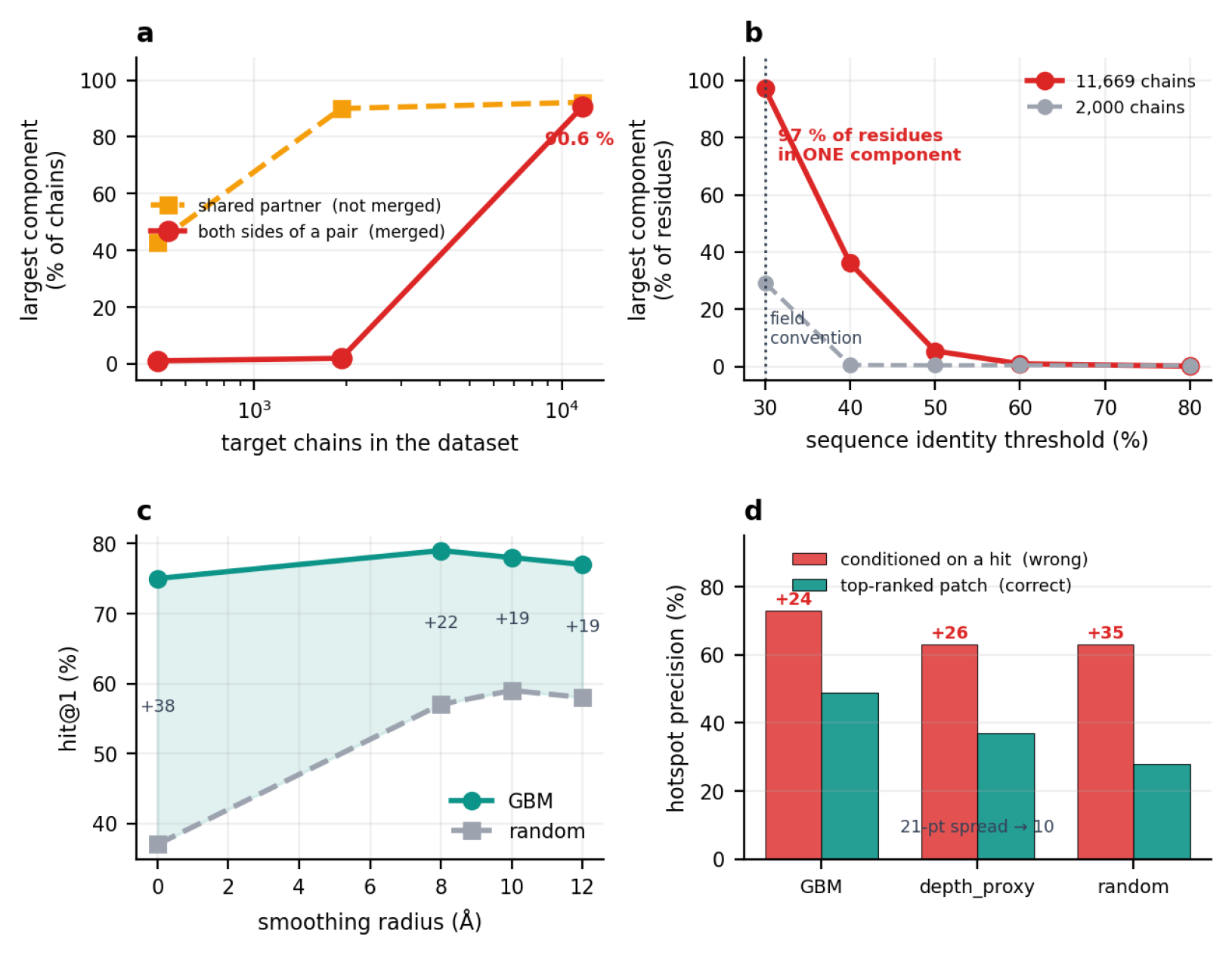
Four ways this task is easy to measure wrong. (a) Both partner graphs percolate with dataset size; the dimer graph, which the split code merges, is negligible at pilot scale and 90.6 % at full scale. (b) Homology single linkage at the conventional 30 % threshold puts 97 % of the complete set in one component, against 29 % for the 2,000-chain pilot. (c) Under a count-only hit rule, smoothing inflates the random baseline faster than the model. (d) Hotspot precision conditioned on success inflates every model and the weakest one most, compressing a 21-point spread to 10.

Two further leaks are specific to partner-agnostic labelling, and earlier drafts of this work conflated them (Figure 4a).

A *dimer edge* joins two chains that are the two sides of one modelled pair — their labels are literally the two faces of a single interface. This graph is sparse at pilot scale (largest component 1.0 % at 485 chains, 1.9 % at 1,918) and percolates abruptly on the complete set: 26,524 edges, 2.27 per chain, largest component **90.6 %**. A split that is demonstrably safe at pilot scale is worthless at production scale, and nothing short of measuring it would reveal that.

A *shared-partner edge* joins two chains that bind the same third protein, so their positive labels are induced by the same partner. This graph is already percolated in the smallest pilot — largest component 42.9 % at 485 chains, 90.0 % at 1,918, 92.2 % on the complete set. **Our split code merges dimer edges but not shared-partner edges**, so this leak is present in the numbers we report. We could not remove it: with 267,164 such edges over 11,669 chains, driven by hub partners that bind hundreds of targets, enforcing it would collapse the dataset into a handful of folds.

We address the two leaks we can with one mechanism: union-find merging over homology and dimer edges with a cap of 5 % of the dataset per component, processing candidate merges in descending order of identity so the strongest relationships are honoured first and only marginal ones are refused. On the complete set this yields 1,345 independent components (against 11,669 if chains were grouped individually), largest 679 chains, which partition into five folds of ≈294,200 residues plus a held-out set of 1,698 chains that is never trained or tuned on.

#### The cap does not eliminate leakage; it trades leakage for balance

The run log records the trade explicitly:

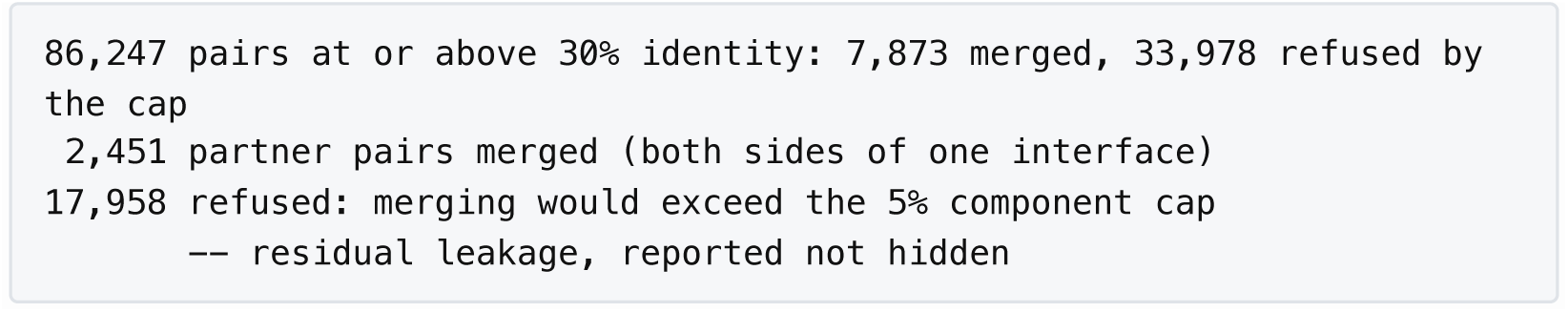

Roughly fifty thousand related pairs are therefore split across folds rather than merged, and the shared-partner graph is not controlled at all. Single-linkage correctness and fold balance cannot both be satisfied on this dataset: enforcing the former puts 97 % of the residues in one component and leaves nothing to cross-validate over, while enforcing the latter leaves measurable residual leakage. We chose balance and report the residue.

Rather than leave that as an unbounded caveat, we measure it and remove it. A chain is marked **partner-contaminated** when a merge involving it was refused by the cap *and* it landed in a different fold from that refused relative — that is, exactly the chains carrying residual leakage. There are 5,813 of them, 62 % of the evaluable set. They remain in training, where leakage costs nothing, and are excluded from every metric.

The bias is real and in the expected direction:

| **subset** | **n** | **hit@1** | **hit@3** | **patch precision** |
|---|---|---|---|---|
| partner-contaminated | 5,813 | 66 % | 81 % | 59 % |
| clean | 3,529 | 63 % | 78 % | 55 % |

Contamination inflates hit@1 by **3.1 points**. Every number in this paper is computed on the clean subset. The residue-level AUC moves the other way — 0.691 pooled before the filter, 0.719 after — because the filter preferentially removes hub chains, whose unions over many partners label much of the surface positive and are correspondingly harder to rank. The two metrics are measuring different things on different populations, and neither direction should be read as the filter “helping” or “hurting”. We note that this correction is only possible because the split code records which merges it refused; a pipeline that caps component size without logging the refusals has the same bias and no way to measure it.

The wider point stands: on a dataset this dense, a benchmark reporting clean homology-controlled splits has either used a stricter threshold than 30 %, used complete rather than single linkage, or not measured its component sizes.

We report all of this because the unguarded version of either clustering gives materially better numbers, and nothing in the output signals that anything is wrong.

## 2.6 Per-target instability

The most consequential result in this paper is the one that is invisible in any aggregate table.

We trained the identical pipeline on three nested training sets — 500, 2,000 and 11,669 chains — and compared the predictions each makes on the chains all pairs have in common.

The best-powered comparison is the middle row: over 277 shared chains the aggregate hit@1 moves 2 points while **36 individual chains — 13 % — change answer**, split in both directions (Figure 5a). Across all three comparisons 13–21 % of chains flip, and per-chain precision correlates at only r ≈ 0.5–0.7 between runs.

**Figure 5.**
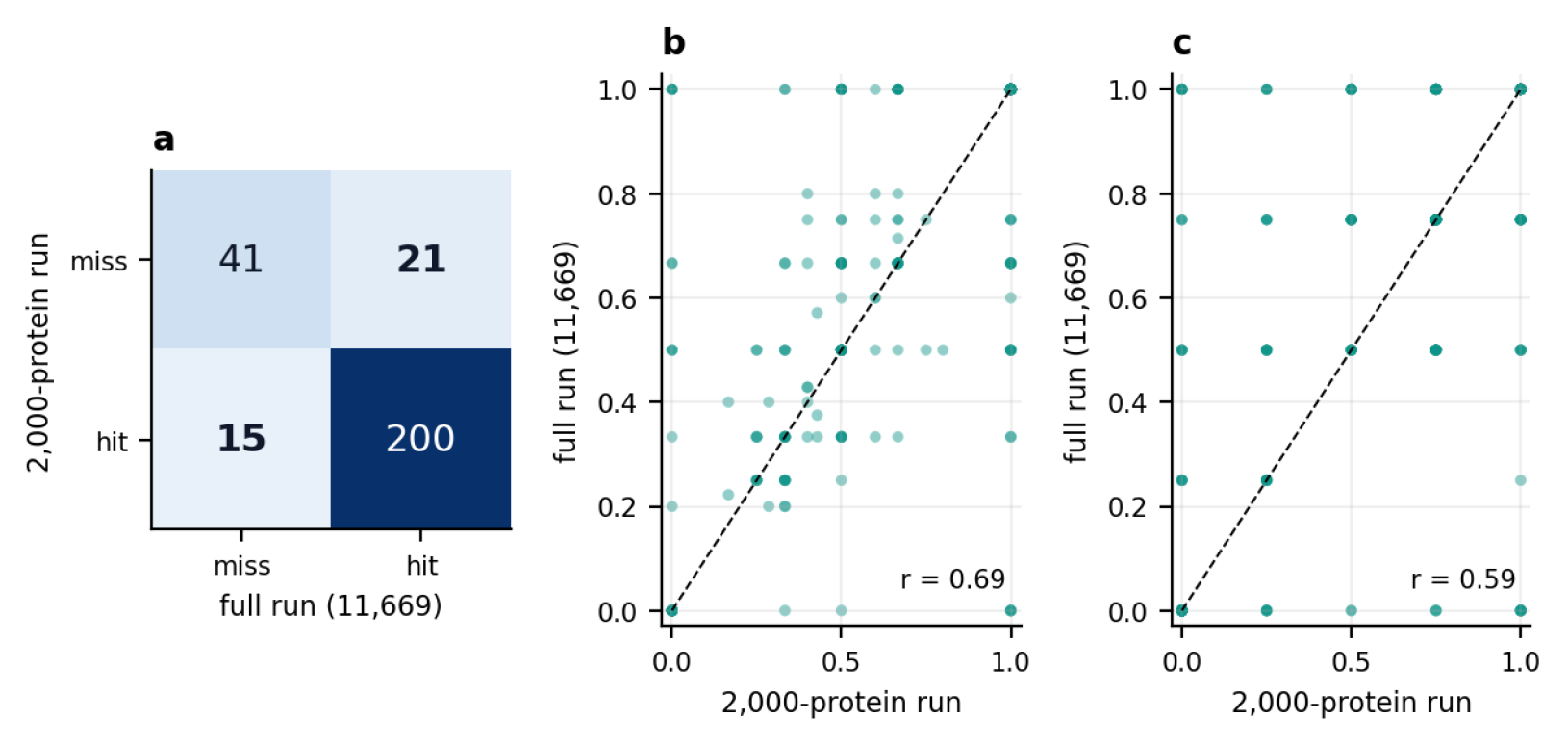
The aggregate is reproducible; individual chains are not. (a) hit@1 agreement between the 2,000-chain and full models over the 277 uncontaminated chains they share: the aggregate moves 2 points while 13 % of chains change answer. (b, c) Perchain patch and hotspot precision between the two runs.

The aggregate is not always stable either: between the 500- and 11,669-chain models it moves 10 points on the 95 chains they share. What is consistent is that the per-chain disagreement is always several times larger than the aggregate difference, and that it does not shrink as the training set grows.

The aggregate is reproducible to within a few points; the individual prediction is not. For a designer with one target and a months-long campaign, the 64 % is close to meaningless and the 13–21 % is the operative risk. We encountered this the hard way: both of our worked examples, IL-12α and IL-6, were written up as successes under the 500-chain model and became misses under the larger one — IL-12α’s top patch goes from 56 % overlap with a true epitope to 16 %, below the hit threshold. We report this rather than re-selecting the examples, and we have replaced the worked-example figure with this measurement.

### 2.7 What did not work

Three substantial efforts produced no improvement and are reported to save others the time.

**Table 3.** Negative results.

| Approach | Result | Cost |
|---|---|---|
| Message-passing GNN with rotation-invariant local-frame edge features | +0.012 AUC, and only on 51 % of chains — indistinguishable from noise | 42× the training time of the GBM |
| Graded regression on ΔSASA instead of binary classification | 0.610 vs 0.624 — worse | Full re-derivation of labels |
| Scaling 500 → 2,000 → 11,669 chains | the lift rose (1.29× → 1.35× → 1.57×) but the headline rate did not improve | 23× the compute |

**Table 4.** Per-chain stability between training sets. All three runs are contamination-filtered, so the comparison is like for like; the shared sets are correspondingly small.

| **Comparison** | **shared chains** | **hit@1** | **flipped** | ***r* (patch prec.)** | ***r* (hotspot)** |
|---|---|---|---|---|---|
| 500 vs 2,000 | 123 | 66 % → 69 % | **21 %** | 0.70 | 0.65 |
| 2,000 vs full | 277 | 78 % → 80 % | **13 %** | 0.69 | 0.59 |
| 500 vs full | 95 | 67 % → 77 % | **18 %** | 0.74 | 0.53 |

The GNN result is the one we expected to go the other way. Geometric deeplearning is the established approach for this task, and a local-frame message-passing network is the natural way to learn 3D residue relationships. It did notbeat gradient boosting on 170 hand-built features here. We attribute this to labelnoise rather than architecture: with partner-agnostic unions over predictedstructures, an unknown share of the negatives are false, and extra modelcapacity fits that noise.

## 3. Discussion

The predictor we built does not beat published interface models and we do not present it as a competitor. What it provides is a controlled setting in which to ask how much of a reported number is signal.

Three recommendations follow for anyone reporting interface or epitope prediction.

### Report lift over a geometry-matched random baseline

A compact random patch hits a real epitope 41 % of the time on this benchmark. Any hit rate reported without that floor is uninterpretable — and the floor is not a constant: it was 52 % on our 2,000-chain pilot, so the same model’s apparent quality shifts by 22 points of lift purely with the composition of the test set. The floor is high because proteins have several binding sites and because epitopes are large; it is a property of the task, not of the evaluation being lax.

### Report patch precision, not just hit rate

Hit@1 asks whether the best guess is right. Patch precision asks what fraction of everything the model emits is right — 46 % here against a 28 % floor. For a user who will test several candidates, that is the number that predicts the cost.

### Report per-target stability

Train two models on different data samples and report how often individual predictions disagree. Across three nested training sets we found 13–21 % of chains changing answer, always several times larger than the aggregate difference between the same two models. We are not aware of an interface-prediction paper that reports this, and we suspect the number is not small for other methods either.

The limitations of this work are substantial and mostly structural. HumanPPI contains predicted, not experimental, complexes; a model trained on it learns “AlphaFold would place a human interface here”, which is correlated with, but not identical to, biological binding. Negatives are unverified. The benchmark is human-only. And because our positives are unions over observed partners, a prediction scored as a false positive may be a genuine binding site for an unmodelled partner — meaning our precision figures are lower bounds, with the inconvenient property that we cannot say by how much.

For the design application that motivated this work, the practical conclusion is narrow and we think defensible: given that de novo binders can now be designed from the target structure alone [15], a single-chain surface model is worth running as a ranking step, because 1.74× enrichment of real contact residues in the top hotspots is a real saving over picking a patch by eye, and it is worth testing two or three patches rather than one, because the gap between hit@1 and hit@3 is 15 points. It is not worth treating the top patch as an answer, and a target whose prediction matters should be scored by more than one model.

## 4. Four ways to measure this task wrong

Each of the following was encountered during development, each produced a plausible result, and each inverted a conclusion we had already recorded. They are collected here because we expect them to recur.

### 4.1 Feature leakage through a complex-derived quantity

`delta_rsa` gave a pooled AUC of 0.999. It is the label as a float. Any quantity computed on the complex — accessibility change, buried area, contact counts — must live in the label file and never in the feature table. The failure is loud when it is total and silent when it is partial.

### 4.2 Percolation in split construction

Covered in §2.5. On the complete set, all three relationship graphs form giant components under single linkage: homology at the conventional 30 % threshold (97.1 % of residues), dimer edges (90.6 % of chains) and shared-partner edges (92.2 %). Two of the three look harmless at pilot scale — the dimer graph holds 1.0 % at 485 chains — so a split validated on a pilot provides no assurance at production scale. The diagnostic is cheap and we recommend it unconditionally: print the size of the largest component as a fraction of the dataset before trusting any split, at the scale you actually intend to train on.

### 4.3 A hit rule that rewards smoothing instead of signal

Our first hit rule was “the patch shares ≥ 3 residues with an epitope”. Under that rule, increasing the smoothing radius from 0 to 12 Å raised random’s hit@1 from 39 % to 62 % (Figure 4c), closing the gap with the model from 37 to 17 points — and at 12 Å the naive depth proxy appeared to beat the trained model. Nothing had been learned; larger patches simply intersect more things. Adding the requirement that ≥ 25 % of the patch lie inside the epitope removes the degeneracy, because a patch cannot grow its way to precision. We had already reported the smoothing result as an improvement before catching this.

### 4.4 Conditioning a quality metric on success

Hotspot precision computed over *successful* patches only, against the same metric computed over the top-ranked patch regardless of whether it hit (Figure 4d):

|  | **conditioned (wrong)** | **top-ranked (correct)** | **inflation** |
|---|---|---|---|
| GBM | 73 % | 49 % | +24 |
| depth proxy | 63 % | 37 % | +26 |
| random | 63 % | 28 % | **+35** |

Conditioning inflates every model, and inflates the worst one most: random succeeds rarely, and when it does it is because it landed squarely on a large epitope, so its hotspots are trivially correct. The effect compresses a 21-point spread between the three methods into a 10-point one, which is enough to make a weak model look competitive.

On the 2,000-chain pilot the same calculation did not merely compress the ranking, it **reversed** it: GBM 77 %, depth proxy 60 %, random 78 %. An earlier draft of this work reported that inversion as a general property of the metric. It is not — it is what this artifact happens to do on that subset. The artifact is general; its severity is dataset-dependent, and we would not have known that without computing it on both. Any metric conditioned on the model being right will flatter a model that is rarely right, by an amount no one can predict in advance.

## 5. Methods

### 5.1 Data

HumanPPI structural models of human protein pairs (AlphaFold2 [10], AlphaFold-Multimer [11], RoseTTAFold [12]), 29,246 pairs, PDB coordinates only, yielding 12,296 target chains and 4,180,280 residues. The main analysis uses the complete set; 500- and 2,000-chain subsets were retained for the stability comparison in §2.6. No external structures were fetched and no structure prediction was run for this work.

### 5.2 Labels

A residue of the target chain is positive if any of its heavy atoms lies within 5.0 Å of any heavy atom of a partner chain, in any complex containing that chain. Labels are the union over all such partners. Solvent accessibility is computed with FreeSASA [16]; relative accessibility uses the Tien et al. theoretical maximum-ASA reference [17]. Distinct epitopes are obtained by agglomerating per-partner contact sets at an overlap coefficient of 0.25.

### 5.3 Features

170 per-residue features computed on the isolated chain only: RSA, burial depth, Cα-shell curvature and convexity, rotation-invariant local-frame neighbourhood descriptors, secondary-structure context, residue identity, standard physicochemical scales, and neighbour means of all of these at 8 Å and 12 Å. No partner-derived quantity, no MSA, no sequence search, no structural alphabet.

### 5.4 Model and splits

`HistGradientBoostingClassifier` (scikit-learn [18]; 400 iterations, learning rate 0.06, 31 leaves, minimum 40 samples per leaf, L2 1.0), a histogram-based gradient booster of the kind introduced by LightGBM [19], with 5-fold cross-validation. Folds are defined over components of a graph whose edges are (i) Smith–Waterman [20] sequence identity ≥ 30 %, computed all-against-all with no k-mer prefilter (Biopython `PairwiseAligner` [21]; no MMseqs2 or Foldseek [7]) and (ii) a shared partner chain. Components are formed by union-find with a cap of 5 % of the dataset, merging in descending order of identity; residual refused merges are counted and reported (§2.5). This gives 1,345 components, five folds of ≈294,200 residues, and a held-out set of 1,698 chains (15 %) that is never trained or tuned on. The smoothing radius is selected per outer fold on the training folds only; it selected 8 Å in all five folds for the model and 10–12 Å for the random baseline. All-against-all alignment over 12,296 chains took 7,070 s on 48 cores.

### 5.5 Evaluation

Patch formation: smooth per-residue scores over a 10 Å Cα neighbourhood, retain the top 25 %, take connected components within 8 Å with a minimum of 5 residues, rank by size × mean score. Hit rule: ≥ 3 shared residues with an epitope **and** ≥ 25 % of the patch inside it. The random baseline places a compact patch of matched size at a random surface position and passes it through the identical pipeline. Hotspot precision is the fraction of the top 4 residues of the top-ranked patch that are true contacts, scored over all chains irrespective of whether the patch hit.

### 5.6 Code and data availability

The evaluation harness, feature extraction, split construction and inference script are available at https://github.com/hello2632/epitope-predictor. HumanPPI is available from its original authors. Inference on a new single-chain structure:

~~~
python scripts/predict.py target.pdb --model model_final.joblib
~~~

`predict.py` calls the training feature scripts as subprocesses rather than reimplementing them, and aborts if the feature columns do not match those the model was trained on — a second implementation drifts, and the model will return confident scores from the wrong inputs.

## Author contributions

B.L. and D.L. jointly conceived the study. B.L. prepared and curated the HumanPPI structural dataset. B.L. and D.L. jointly implemented the feature extraction, split construction, model training and evaluation code, and jointly performed the analysis. B.L. produced the figures. B.L. and D.L. jointly wrote the original draft, and B.L. revised the manuscript. Both authors read and approved the final version and accept responsibility for all of its content.

## Use of generative AI

Claude (Anthropic) was used as a coding and writing assistant: it contributed to the implementation of the analysis scripts and to drafting and revising the text of this manuscript. In keeping with ICMJE and COPE guidance, the tool is not listed as an author, since it cannot take responsibility for the work or approve the submitted version. All code it produced was reviewed by the authors, all analyses were executed and checked by the authors, and the authors take full responsibility for the accuracy and integrity of every result and statement reported here.

## References

1. Watson, J. L. et al. De novo design of protein structure and function with RFdiffusion. Nature 620, 1089–1100 (2023).

2. Pacesa, M. et al. BindCraft: one-shot design of functional protein binders. Nature (2025).

3. Gainza, P. et al. Deciphering interaction fingerprints from protein molecular surfaces using geometric deep learning. Nat. Methods 17, 184–192 (2020).

4. Tubiana, J., Schneidman-Duhovny, D. & Wolfson, H. J. ScanNet: an interpretable geometric deep learning model for structure-based protein binding site prediction. Nat. Methods 19, 730–739 (2022).

5. Krapp, L. F., Abriata, L. A., Cortés Rodriguez, F. & Dal Peraro, M. PeSTo: parameter-free geometric deep learning for accurate prediction of protein binding interfaces. Nat. Commun. 14, 2175 (2023).

6. Foldseek-Interface reveals a protein interface universe far from complete. bioRxiv (2025).

7. van Kempen, M. et al. Fast and accurate protein structure search with Foldseek. Nat. Biotechnol. 42, 243–246 (2024).

8. Jones, S. & Thornton, J. M. Analysis of protein-protein interaction sites using surface patches. J. Mol. Biol. 272, 121–132 (1997).

9. Levy, E. D. A simple definition of structural regions in proteins and its use in analyzing interface evolution. J. Mol. Biol. 403, 660–670 (2010).

10. Jumper, J. et al. Highly accurate protein structure prediction with AlphaFold. Nature 596, 583–589 (2021).

11. Evans, R. et al. Protein complex prediction with AlphaFold-Multimer. bioRxiv (2022).

12. Baek, M. et al. Accurate prediction of protein structures and interactions using a three-track neural network. Science 373, 871–876 (2021).

13. Burke, D. F. et al. Towards a structurally resolved human protein interaction network. Nat. Struct. Mol. Biol. 30, 216–225 (2023).

14. Humphreys, I. R. et al. Computed structures of core eukaryotic protein complexes. Science 374, eabm4805 (2021).

15. Cao, L. et al. Design of protein-binding proteins from the target structure alone. Nature 605, 551–560 (2022).

16. Mitternacht, S. FreeSASA: an open source C library for solvent accessible surface area calculations. F1000Research 5, 189 (2016).

17. Tien, M. Z., Meyer, A. G., Sydykova, D. K., Spielman, S. J. & Wilke, C. O. Maximum allowed solvent accessibilities of residues in proteins. PLoS ONE 8, e80635 (2013).

18. Pedregosa, F. et al. Scikit-learn: machine learning in Python. J. Mach. Learn. Res. 12, 2825–2830 (2011).

19. Ke, G. et al. LightGBM: a highly efficient gradient boosting decision tree. NeurIPS 30 (2017).

20. Smith, T. F. & Waterman, M. S. Identification of common molecular subsequences. J. Mol. Biol. 147, 195–197 (1981).

21. Cock, P. J. A. et al. Biopython: freely available Python tools for computational molecular biology and bioinformatics. Bioinformatics 25, 1422–1423 (2009).

